# Comparative evaluation of sample preparation and enrichment methods for culture and molecular detection of *Burkholderia pseudomallei* in naturally contaminated soils from Thailand over one year

**DOI:** 10.64898/2026.10.26.761812

**Authors:** Charlene Mae Salao Cagape, Théo Dokhelar, Natnaree Saiprom, Noppadol Prasertsincharoen, Mégane Gasqué, Karine Laroucau, Narisara Chantratita, Vanina Guernier-Cambert

**Author notes:** These authors contributed equally to this work (Charlene Mae Salao Cagape, Théo Dokhelar).

## Abstract

**Background:** *Burkholderia pseudomallei* is a soil-dwelling bacterium that causes melioidosis. Accurate environmental detection of *B. pseudomallei* is critical for surveillance strategies and for tracing the source of human infection. However, the detection of *B. pseudomallei* in soil samples, which are complex matrices with a high microbial background, remains difficult. This study aimed to compare the culture and the molecular detection of *B. pseudomallei* with or without pre-enrichment in naturally contaminated soils from Thailand.

**Methodology/principal findings:** In 2025, a total of 640 soil samples were collected from 10 sites from different environments and sampling periods in Amnat Charoen Province. The soils were evaluated for the presence of *B. pseudomallei* either without enrichment or following enrichment in Ashdown or ACER broths, using culture and real-time PCR detection methods. qPCR consistently identified a higher number of *B. pseudomallei* positive samples than culture regardless of the pre-treatment method and sampling period. ACER enrichment nearly doubled the odds of detection compared with no enrichment (OR = 1.90) or with Ashdown enrichment (OR = 2.09), with a model-estimated mean Ct difference of 13.6 between ACER enrichment and no enrichment. Overall, qPCR yielded higher detection rates than culture for *B. pseudomallei* in soil samples, particularly following enrichment.

**Conclusions/significance:** This study highlights the importance of pre-enrichment protocols with detection techniques to successfully identify *B. pseudomallei* in environmental samples. For both culture and molecular detection methods, *B. pseudomallei* was better detected when the soil samples were enriched in ACER broth compared to Ashdown broth and without enrichment. These findings highlight the need for combining optimal selective enrichment strategies with detection techniques for environmental surveillance. Although ACER enrichment improved the recovery of viable strains, culture recovered *B. pseudomallei* from only a fraction of qPCR-positive samples, highlighting the need for more selective media to facilitate isolation of *B. pseudomallei*.

## Introduction

*Burkholderia pseudomallei* is a gram-negative and facultative intracellular bacterium that resides in the rhizosphere of tropical soils and causes melioidosis, a disease that claims an estimated 89,000 lives annually [1,2]. The infection occurs through direct contact with contaminated soil or water, primarily via percutaneous inoculation, inhalation, or ingestion [1]. Melioidosis is endemic in Southeast Asia and northern Australia, with Thailand considered one of the major endemic hotspots worldwide [2]. In Thailand, the disease represents an important public health concern, particularly among rice farmers and agricultural workers as well as populations exposed to wet soils and surface water during the rainy season or during flooding [3]. Environmental investigations aiming at detecting the bacterium in soil are essential to better understand its ecology, environmental persistence, and transmission pathways. Such environmental studies can contribute to mapping the spatial distribution of *B. pseudomallei* and targeting regions at increased risk for melioidosis. Environmental surveillance may also improve predictive models of disease emergence and support public health prevention strategies in endemic areas. However, *B. pseudomallei* has proven difficult to detect in complex environmental matrices for several reasons: (i) its highly heterogeneous spatial distribution in soil, both horizontally and vertically, which may also vary with the season, making it difficult to predict where the bacterium is likely to be found; (ii) its low concentrations when present; and (iii) the inherent complexity of performing culture or PCR on soil samples (e.g., culture contamination, presence of PCR inhibitors) [4–6]. Additionally, the bacterium may enter dormant or viable-but-non-culturable states [7], although these physiological states remain largely unexplored in this species.

To improve the detection of *B. pseudomallei* in soil samples, the Detection of Environmental *Burkholderia pseudomallei* Working Party (DEBWorP) proposed consensus guidelines for environmental sampling in 2013 [8]. However, the soil culture protocol involving an enrichment step in threonine-basal salt solution with colistin (TBSS-C50), followed by subculturing on Ashdown agar, which has long been recommended and included in these consensus guidelines, has been shown to lack sensitivity [9,10].

Hence, melioidosis researchers have developed new, more selective enrichment media and explored molecular methods as alternatives or complementary approaches to conventional culture. MacConkey agar has been modified to improve the isolation of *B. pseudomallei* from soil by incorporating four antimicrobial agents, i.e., vancomycin, amphotericin B, gentamicin and colistin [11]. TBSS-C50 has also been modified by using erythritol as a carbon source [9], and was more recently further improved into a phosphate-buffered acidic erythritol medium (ACER) [12]. While several comparison studies of detection methods have been published, many have relied on spiked soil samples, i.e., soils experimentally inoculated with known concentrations or dilutions of pure *B. pseudomallei* cultures [10]. Such approaches are essential for assessing the sensitivity, specificity, and limit of detection of different methods under controlled conditions; however, they may not accurately reflect the complexity of naturally contaminated soils collected in endemic areas. This limitation may be even more pronounced when spiked soils have been autoclaved or otherwise sterilized prior to inoculation, as the absence of the native soil microbiota and natural organic compounds may affect bacterial behavior and impact diagnostic performance. The objective of this study was therefore to assess which method or combination of methods provide the most effective detection of *B. pseudomallei* in naturally contaminated soils from different sites and sampling seasons in Thailand.

## Material and method

### Ethics statement

The sampling sites included both public areas (forests) and private properties (crop fields, gardens). Oral informed consent was obtained from all farmers and landowners prior to sample collection. Soil sampling was conducted in collaboration with the regional office of the Land Development Department (LDD) in Amnat Charoen, Thailand. The study was approved by the Mahidol University Institutional Biosafety Committee (MU-IBC 2024–013). All experiments involving *B. pseudomallei* were performed in an enhanced biosafety level 2 (BSL2+) laboratory. All experiments were performed in accordance with relevant guidelines and regulations.

### Soil sampling

During 2025, four soil sampling campaigns were conducted in Mueang Amnat Charoen district, Amnat Charoen Province, in northeastern Thailand: 12 February (dry season), 27-28 May (early wet season), 19 August (wet season), and 30-31 October (early dry season). Ten sites from various environments were selected within a 4-km radius. At each site, 16 soil samples were collected at 2-m intervals following a 6 x 6m square grid, at a depth of 40 cm using a steel auger. After digging the primary sampling holes, individual metal spoons disinfected with 70% ethanol were used to remove the first layer of soil potentially contaminated by the auger and to collect 100 g of soil into single-use plastic zip bags.

Soil samples were stored at ambient temperature in the dark and transferred to the laboratory at the Faculty of Tropical Medicine, Mahidol University, Bangkok, within three days of collection, and processed immediately. Due to a several-months closure of the BSL2+ laboratory at the Faculty of Tropical Medicine, Mahidol University, Bangkok, samples collected in October 2025 (400 g rather than the usual 100 g to help maintain sample integrity during prolonged storage) were double-bagged and stored in the dark in Styrofoam boxes in a laboratory room maintained at 26°C until processing in January 2026.

### Laboratory procedures

We compared the direct detection (by culture) and the molecular detection (by real-time PCR) of *B. pseudomallei* in all soil samples following three pre-treatment protocols: no enrichment, Ashdown enrichment, or ACER enrichment.

#### Enrichment, culture, and screening of suspected colonies

First, 100 mL of distilled water was added directly to 100 g of soil in each zip bag, and the mixture was thoroughly homogenized manually until soil aggregates were dispersed. Samples were then left to settle overnight. The following day, 3.2 mL of the upper layer of this mixture (hereafter referred to as the soil supernatant) was collected in two microtubes and used as follows: 10 µL and 100 µL were directly plated onto Ashdown agar [13] and incubated at 42°C for four days; 1 mL was used for Ashdown enrichment (prior to agar plating); 1 mL was used for ACER enrichment (prior to agar plating); 1 mL was used directly for DNA extraction and molecular testing (see next section) (Fig 1). After two days of enrichment in Ashdown broth [14] and nine days in ACER broth [10,12], 10 µL of each enrichment culture were plated onto Ashdown agar and incubated at 42°C for four days; 1 mL of each enrichment culture was used for DNA extraction and molecular testing (see next section) (Fig 1).

**Fig 1.**
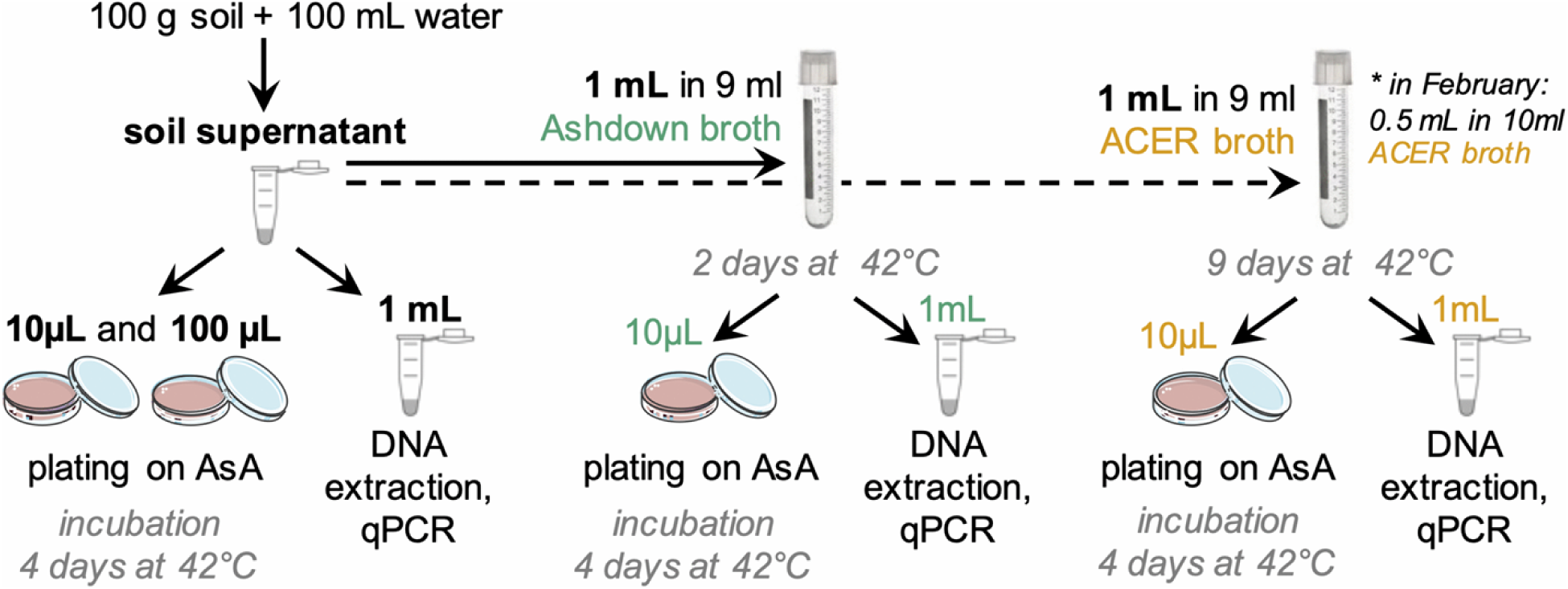
Flowchart of the laboratory procedures comparing three sample preparation methods for the detection of *B. pseudomallei* in soil samples. Soil supernatants were subjected to either no enrichment, or Ashdown enrichment, or ACER enrichment prior to detection by culture and real-time PCR (qPCR). For qPCR, DNA was extracted from pellets obtained either directly (no enrichment) or after enrichment. AsA: Ashdown agar.

In February, ACER enrichment was performed using only 0.5 mL of soil supernatant in 10 mL of ACER broth, following a previously published protocol [10], whereas for the three subsequent sampling periods, the protocol was adjusted to 1 mL of soil supernatant in 9 mL of ACER broth to match the dilution ratio used for the Ashdown enrichment procedure.

Suspected *B. pseudomallei* colonies on Ashdown agar were confirmed by latex agglutination [15] and enumerated. All plates with at least one agglutinating colony were considered culture-positive for comparison with the molecular results.

#### DNA extraction and molecular testing

Of the 3.2 mL of soil supernatant previously collected, 1 mL was centrifuged at 8,000 rpm for 10 min and the resulting supernatant was discarded. Following enrichment, 1 mL of each enrichment broth was centrifuged in the same way and the supernatant discarded. DNA was extracted from the resulting pellets using the DNeasy PowerSoil Pro Kit (QIAGEN, Hilden, Germany), following the manufacturer’s instructions.

All extracted gDNA samples were tested by qPCR using primers and a probe that target a 115 bp fragment of the TTS1-*orf2* gene from *B. pseudomallei* type III secretion system: BpTT4176F (5′-CGTCTCTATACTGTCGAGCAATCG-3′), BpTT4290R (5′-CGTGCACACCGGTCAGTATC-3′), BpTT4208P (5′-CCGGAATCTGGATCACCACCACTTTCC-3′) [16]. Molecular assays were performed on a Bio-Rad CFX Opus 96 system using the iTaq Universal Probes One-Step kit (Bio-Rad). While a reverse transcription step can increase bacterial detection sensitivity, the DNeasy PowerSoil Pro Kit is designed for DNA extraction, with no RNA co-purification step, so reverse transcription did not contribute to detection. Each reaction contained 3 μL of DNA template and 17 μL of reaction mix including 1X iTaq Universal Probes reaction mix, 1X of iScript Reverse Transcriptase, 0.4 μM of each primer and 0.2 μM of probe. Sterile water was added to bring the final volume to 20 µL. The amplification programme consisted of an initial incubation at 50°C for 10 min, followed by denaturation at 95°C for 3 min, then 45 cycles of 95°C for 15 sec and 60°C for 30 sec. Samples yielding a quantification cycle (Ct) value > 38 were retested. Samples tested in duplicate and showing discordant results (one negative and one positive) were tested a third time. For Ct-specific analyses, qPCR-negative samples were excluded; for qPCR-positive samples with multiple Ct values, a mean Ct value was calculated for each individual pre-treatment protocol. For samples collected from sites 1 to 5 during the October sampling period, no 1 mL aliquot of Ashdown enrichment broth was collected for subsequent analysis; qPCR after Ashdown enrichment was performed only from sites 6 to 10. As a result, molecular comparisons were based on 160 samples in February, May, August, but were limited to 80 samples for the October sampling period.

### Statistical analysis

#### Agreement and comparison of molecular results across three pre-treatment protocols

For each soil sample, *B. pseudomallei* detection status was coded as a binary variable (0 = negative, 1 = positive) following qPCR with different pre-treatment protocols. The three protocols were compared pairwise (three pairwise comparisons in total). For each pair, agreement was assessed using Cohen’s kappa, and directional asymmetry in discordant pairs was assessed using McNemar’s test, both overall and stratified by sampling season. Kappa coefficients were interpreted following standard thresholds, ranging from poor (κ ≤ 0) to near perfect agreement (κ = 0.81–1.0) [17]. A significant McNemar test (p < 0.05) indicated asymmetry in discordant pairs, meaning one pre-treatment protocol systematically detected more positives than the other.

For the October sampling period, analyses were restricted to samples collected from sites 6 to 10, for which qPCR results from all three pre-treatment protocols were available (n = 80). Consequently, qPCR−based agreement metrics and model estimates for this season were based on this subset only. Euler diagrams were used to visualize the overlap in molecular diagnosis results among the three pre-treatment protocols.

#### Modeling of detection probability between molecular detection methods x pre-treatment protocols

To account for the hierarchical structure of the dataset, including repeated measurements of sampling sites across four seasons, a generalized linear mixed model (GLMM) with a binomial distribution and logit link function was fitted, with *B. pseudomallei* detection status (presence / absence) as the binary response variable. In this model, we assessed the effect of the pre-treatment protocol (three levels: no enrichment, Ashdown enrichment, and ACER enrichment) and the detection method (two levels: culture and qPCR). *B. pseudomallei* detection probabilities were compared between the three pre-treatment protocols within each detection method, and between qPCR and culture within each pre-treatment protocol, with detection status coded separately for each sample x pre-treatment x detection method combination:

*B. pseudomallei detection status* ∼ *pre-treatment* * *method* + *(1* | *Sample)* + *(1* | *Site)* + *(1* | *Season)*

In the model, pre-treatment protocol, detection method, and their interaction were included as fixed effects, whereas sample, site, and season were included as random intercepts to account for the non-independence of repeated observations from the same sampling point as well as for possible variation between sites and sampling periods. Estimated marginal means (EMMs) and pairwise contrasts, with Tukey-adjusted p-values and 95% confidence intervals, were used to compare detection probabilities.

#### Modeling of qPCR Ct value change between three pre-treatment protocols

To compare qPCR performance for the three pre-treatment protocols (no enrichment, Ashdown enrichment or ACER enrichment) while accounting for repeated measurements across four seasons, Ct values from samples detected by qPCR following all three pre-treatment protocols were analyzed using a Gaussian GLMM:

*B. pseudomallei Ct value ∼ pre-treatment* + *(1* | *Season)*

This paired analysis, restricted to samples that were qPCR-positive under all three protocols, reduced potential bias associated with missing Ct values resulting from qPCR-negative results. Because each sample contributed a Ct value under all three protocols, protocol effects were estimated within samples. Sample- and site-level random intercepts had estimated variances of zero and were therefore not retained; sampling season was included as a random intercept. Pairwise comparisons of Ct values between pre-treatment protocols were performed using EMMs with Tukey adjustment for multiple testing. Spearman rank correlation analysis was used to assess whether pre-treatment protocols preserved the relative ranking of target DNA abundance based on Ct values among samples that were qPCR-positive under all three pre-treatment protocols. Two-tailed *p*-values < 0.05 were considered statistically significant.

Because a different supernatant-to-broth ratio was used for ACER enrichment in February, analyses were repeated excluding the February sampling period as a sensitivity analysis. All analyses were performed in R version 4.5.2 [18] using glmmTMB [19], emmeans [20] and caret [21] packages. Model fit was assessed for singularity and overdispersion, with overdispersion evaluated using simulated residuals from the DHARMa package [22].

## Results

### Detection of B. pseudomallei in soil samples

Overall, 160 samples (16 samples from each of the 10 sites) were collected during each sampling period, resulting in a total of 640 samples. On Ashdown agar plates, *B. pseudomallei* isolates were difficult to identify due to background contamination; therefore, only a limited number of plates yielded confirmed positive results by agglutination, with some seasonal variation (Table 1). Molecular detection by qPCR consistently identified a higher number of positive soil samples than culture, regardless of the pre-treatment protocol or sampling season (Table 1). Detailed molecular and culture results for individual soil samples following each of the three pre-treatment protocols are provided as Supporting Information (Table S1).

**Table 1.** Summary of culture- and qPCR-positive samples according to the sample pre-treatment protocol and sampling period. In the “culture” column, each soil sample is counted once, whatever the number of *B. pseudomallei*-positive agar plates or the number of *B. pseudomallei* isolates per plate, if any.

| Sampling period | No enrichment |  | Ashdown enrichment |  | ACER enrichment |  |
| --- | --- | --- | --- | --- | --- | --- |
|  | culture | qPCR | culture | qPCR | culture | qPCR |
| February | 3/160 | 41/160 | 0/160 | 40/160 | 15/160<sup>a</sup> | 29/160 |
| May | 0/160 | 20/160 | 0/160 | 12/160 | 3/160<sup>b</sup> | 24/160 |
| August | 0/160 | 20/160 | 0/160 | 25/160 | 1/160<sup>c</sup> | 52/160 |
| October | 0/160 | 52/160 | 8/160 | 28/80<sup>d</sup> | 20/160 | 94/160 |
| Total | 3/640 | 133/640 | 8/640 | 105/560 | 39/640 | 199/640 |

<sup>a</sup> 2/15 culture-positive samples were qPCR-negative. <sup>b</sup> 3/3 culture-positive samples were qPCR-negative.
<sup>c</sup> 1/1 culture positive sample was qPCR-positive after no enrichment but qPCR-negative after ACER enrichment. <sup>d</sup> In October, 1 mL of Ashdown broth was collected only from sites 6 to 10 for DNA extraction and qPCR testing.

### Agreement and comparison of qPCR results across three pre-treatment protocols

As isolation on Ashdown agar was limited, comparison analyses focused on molecular detection results obtained using three pre-treatment protocols: no enrichment, Ashdown enrichment, or ACER enrichment (Table 2). Ashdown enrichment showed fair agreement with no enrichment (κ = 0.39), with no significant asymmetry (*p* = 0.627), indicating similar proportions of positive samples detected by the two protocols. ACER enrichment showed moderate agreement with no enrichment (κ = 0.43), with significant asymmetry (*p* < 0.001), indicating that ACER enrichment detected additional positive samples that were not detected without enrichment. Ashdown enrichment and ACER enrichment also showed moderate agreement (κ = 0.46), with significant asymmetry (*p* < 0.001), consistent with the higher overall detection rate observed with ACER enrichment (Table 1).

**Table 2.** Agreement and comparison of *B. pseudomallei* qPCR results across three pre-treatment protocols, overall and by sampling period. Pre-treatment protocols are: no enrichment; Ashdown enrichment; ACER enrichment. Kappa values range from 0 to 1 (*k* ≤ 0: poor agreement; 0.01–0.20: slight agreement; 0.21–0.40: fair agreement; 0.41–0.60: moderate agreement; 0.61–0.80: substantial agreement; 0.81–1.0: almost perfect agreement) [17]. A McNemar test *p*-value < 0.05 indicates a statistically significant difference in paired detection rates between the compared approaches; significant *p*-values are shown in bold.

| Sampling period | Sample pre-treatment protocols | Kappa index | McNemar $p$-value |
| --- | --- | --- | --- |
| Overall | No enrichment vs Ashdown enrichment | 0.39 | 0.627 |
|  | No enrichment vs ACER enrichment | 0.43 | **< 0.001** |
|  | Ashdown enrichment vs ACER enrichment | 0.46 | **< 0.001** |
| February | No enrichment vs Ashdown enrichment | 0.22 | 1.000 |
|  | No enrichment vs ACER enrichment | 0.38 | 0.059 |
|  | Ashdown enrichment vs ACER enrichment | 0.54 | **0.046** |
| May | No enrichment vs Ashdown enrichment | 0.24 | 0.136 |
|  | No enrichment vs ACER enrichment | 0.47 | 0.502 |
|  | Ashdown enrichment vs ACER enrichment | 0.38 | **0.014** |
| August | No enrichment vs Ashdown enrichment | 0.61 | 0.302 |
|  | No enrichment vs ACER enrichment | 0.36 | **< 0.001** |
|  | Ashdown enrichment vs ACER enrichment | 0.46 | **< 0.001** |
| October | No enrichment vs Ashdown enrichment | 0.41 | 0.831 |
|  | No enrichment vs ACER enrichment | 0.40 | **< 0.001** |
|  | Ashdown enrichment vs ACER enrichment | 0.27 | **< 0.001** |

When stratified by sampling period, Ashdown enrichment and no enrichment showed no significant asymmetry in any period (κ = 0.22–0.61; *p* > 0.05 for all comparisons), indicating that their detection rates were comparable across sampling season. ACER enrichment and no enrichment showed significant asymmetry in August and October (both *p* < 0.001), but not in February (*p* = 0.059) or May (*p* = 0.502) (κ = 0.36–0.47), suggesting that the additional detection provided by ACER enrichment over no enrichment was not consistent across sampling periods. Ashdown enrichment and ACER enrichment showed significant asymmetry in all four sampling periods (*p* ≤ 0.046; κ = 0.27–0.54), indicating that ACER enrichment yielded higher detection rates than Ashdown enrichment across all sampling periods.

Euler diagrams illustrated the overlap in positive samples detected by qPCR following the three pre-treatment protocols. Across all sampling periods, ACER enrichment yielded the largest number of unique positive samples (60 samples not detected by either of the other two protocols), compared with no enrichment (35 unique positive samples) and Ashdown enrichment (28 unique positive samples) (Fig 2A). Fifty-four samples were detected by qPCR following all three pre-treatment protocols, representing the core overlap. In addition, 21 samples were detected by both ACER enrichment and no enrichment but not by Ashdown enrichment, 22 by both ACER and Ashdown enrichment but not by no enrichment, and one by Ashdown enrichment and no enrichment but not by ACER enrichment. Detection patterns varied across sampling periods (Fig 2B to 2E).

**Fig 2.**
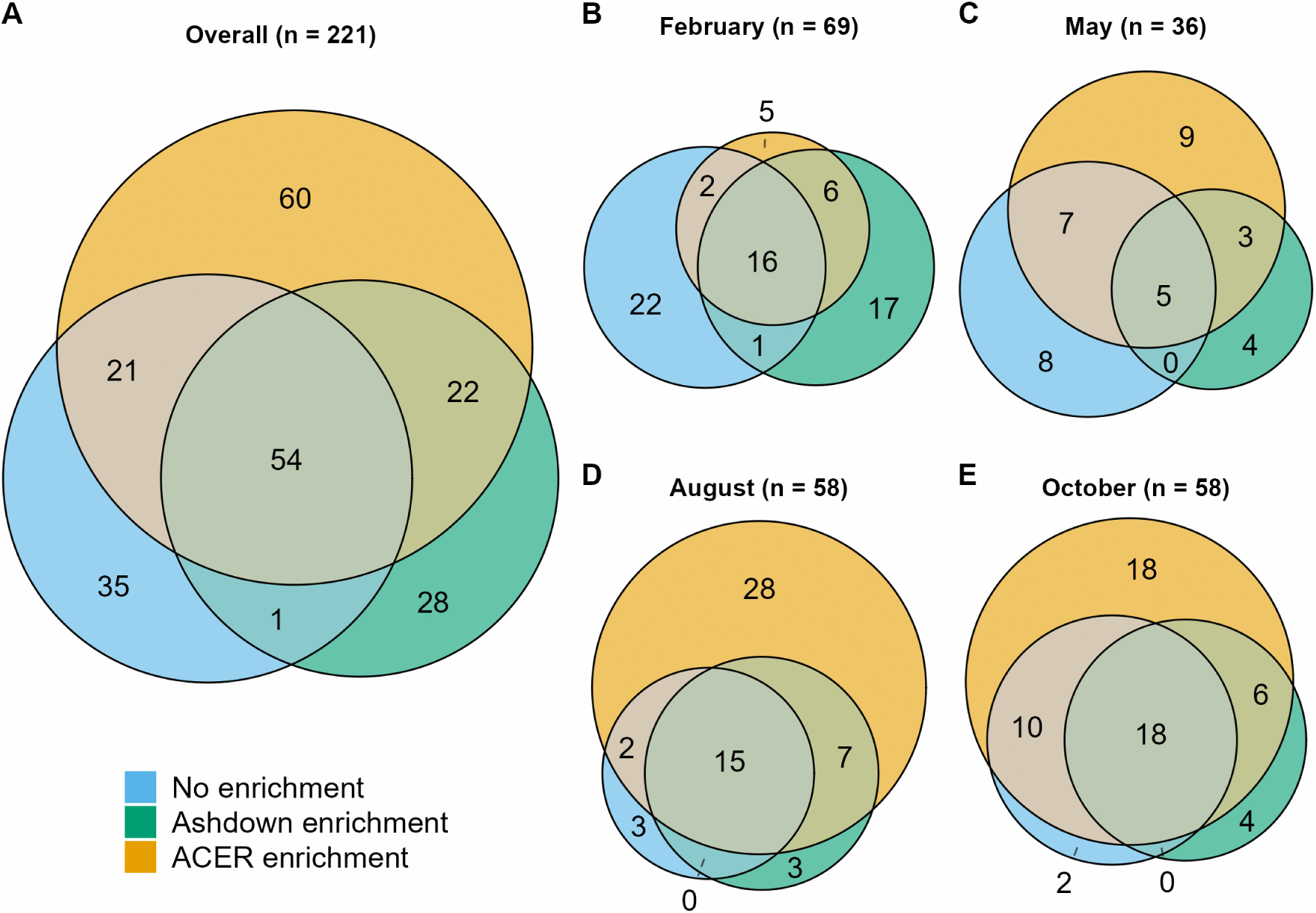
Euler diagrams showing the overlap in *B. pseudomallei*-positive samples detected by qPCR following three sample pre-treatment protocols (no enrichment, Ashdown enrichment, ACER enrichment). Numbers indicate the number of qPCR-positive samples uniquely detected or shared among the three protocols. (A) Overall results (n = 221 positive / 560 tested); (B) February 2025 (n = 69 positive / 160 tested); (C) May 2025 (n = 36 positive / 160 tested); (D) August 2025 (n = 58 positive / 160 tested); (E) October 2025 (n = 58 positive / 80 tested). For October, only samples with available qPCR results for all three pre-treatment protocols (sites 6 −10) were included.

### Modeling the effect of pre-treatment on detection by qPCR and culture

GLMM analysis showed that the pre-treatment protocol significantly affected the probability of *B. pseudomallei* detection, both for qPCR (χ² = 22.81, df = 2, *p* < 0.0001) and for culture (χ² = 36.53, df = 2, *p* < 0.0001) (Fig 3, Table S2).

**Fig 3.**
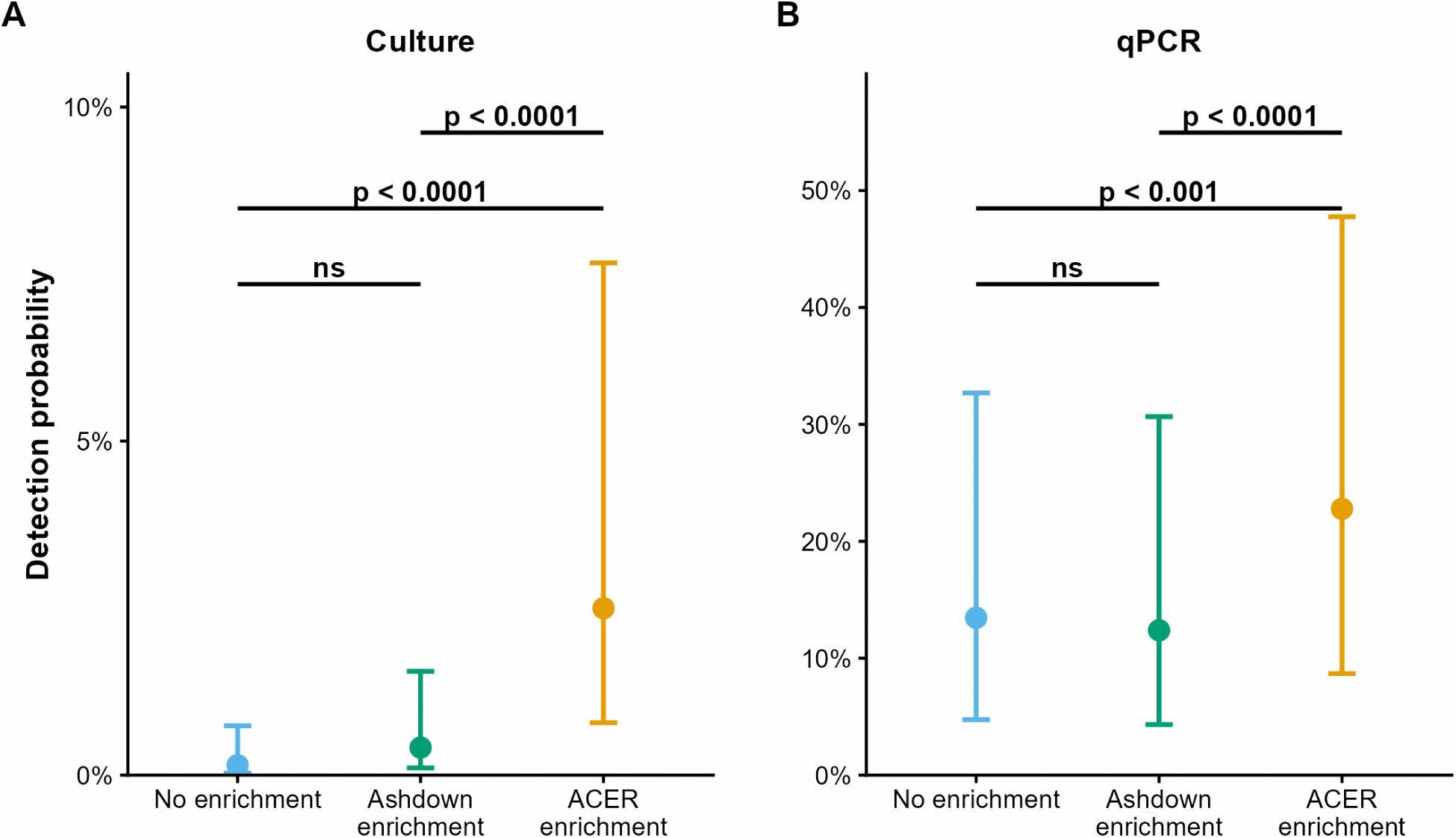
Model-estimated probabilities of *B. pseudomallei* detection following no enrichment, Ashdown enrichment, or ACER enrichment of soil samples, by detection method with (A) qPCR and (B) culture. The y-axis scales differ between panels A and B to accommodate the much lower detection probabilities obtained by culture. Points represent model-predicted *B. pseudomallei* detection probabilities, colored by pre-treatment protocol. Error bars indicate 95% confidence intervals. Brackets show Tukey-adjusted pairwise comparisons between pre-treatment protocols within each detection method (*p*-values).

For qPCR, the model-estimated detection probabilities were 13.5% for no enrichment, 12.4% for Ashdown enrichment, and 22.8% for ACER enrichment (Fig 3A). Based on the model, ACER enrichment was associated with approximately twice the odds of detection compared with no enrichment (OR = 1.90, 95% CI = 1.28–2.82, *p* < 0.001) and Ashdown enrichment (OR = 2.09, 95% CI = 1.40–3.11, *p* < 0.001). Ashdown enrichment did not significantly differ from no enrichment (OR = 0.91, 95% CI = 0.60–1.38, *p* = 0.855).

For culture, the model-estimated detection probabilities were 0.15% for no enrichment, 0.4% for Ashdown enrichment, and 2.5% for ACER enrichment (Fig 3B). Based on the model, ACER enrichment was associated with significantly higher odds of detection than no enrichment (OR = 17.26, 95% CI = 4.10–72.69, *p* < 0.0001) and Ashdown enrichment (OR = 6.19, 95% CI = 2.37–16.14, *p* < 0.0001), while Ashdown enrichment did not significantly differ from no enrichment (OR = 2.79, 95% CI = 0.55–14.05, *p* = 0.298)

The model showed that the difference between qPCR and culture depended on the pre-treatment protocol. With every pre-treatment protocol, the model-estimated odds of *B. pseudomallei* detection were higher by qPCR than by culture, and the magnitude of this difference varied across protocols: it was largest with no enrichment (OR = 104.72, 95% CI = 31.97–342.97, *p* < 0.0001), smaller with Ashdown enrichment (OR = 34.16, 95% CI = 15.73–74.19, *p* < 0.0001), and smallest with ACER enrichment (OR = 11.51, 95% CI = 7.40–17.89, *p* < 0.0001). Compared with no enrichment, the increase in detection odds with ACER enrichment was significant for both qPCR and culture, with a greater increase for culture than for qPCR (*p* < 0.001). In contrast, the corresponding difference between Ashdown enrichment and no enrichment was not significant (*p* = 0.116). No overdispersion or singular fit was detected in this model.

### Distribution of qPCR Ct values between three pre-treatment protocols

While non-enriched qPCR positive samples generally displayed high Ct values, reflecting low amounts of detectable target DNA in soils, both enrichment protocols resulted in substantial Ct reductions, consistent with increased *B. pseudomallei* DNA abundance following enrichment (Fig 4 and Fig S1). However, the magnitude of Ct reduction differed among samples, suggesting heterogeneous bacterial growth during enrichment.

**Fig 4.**
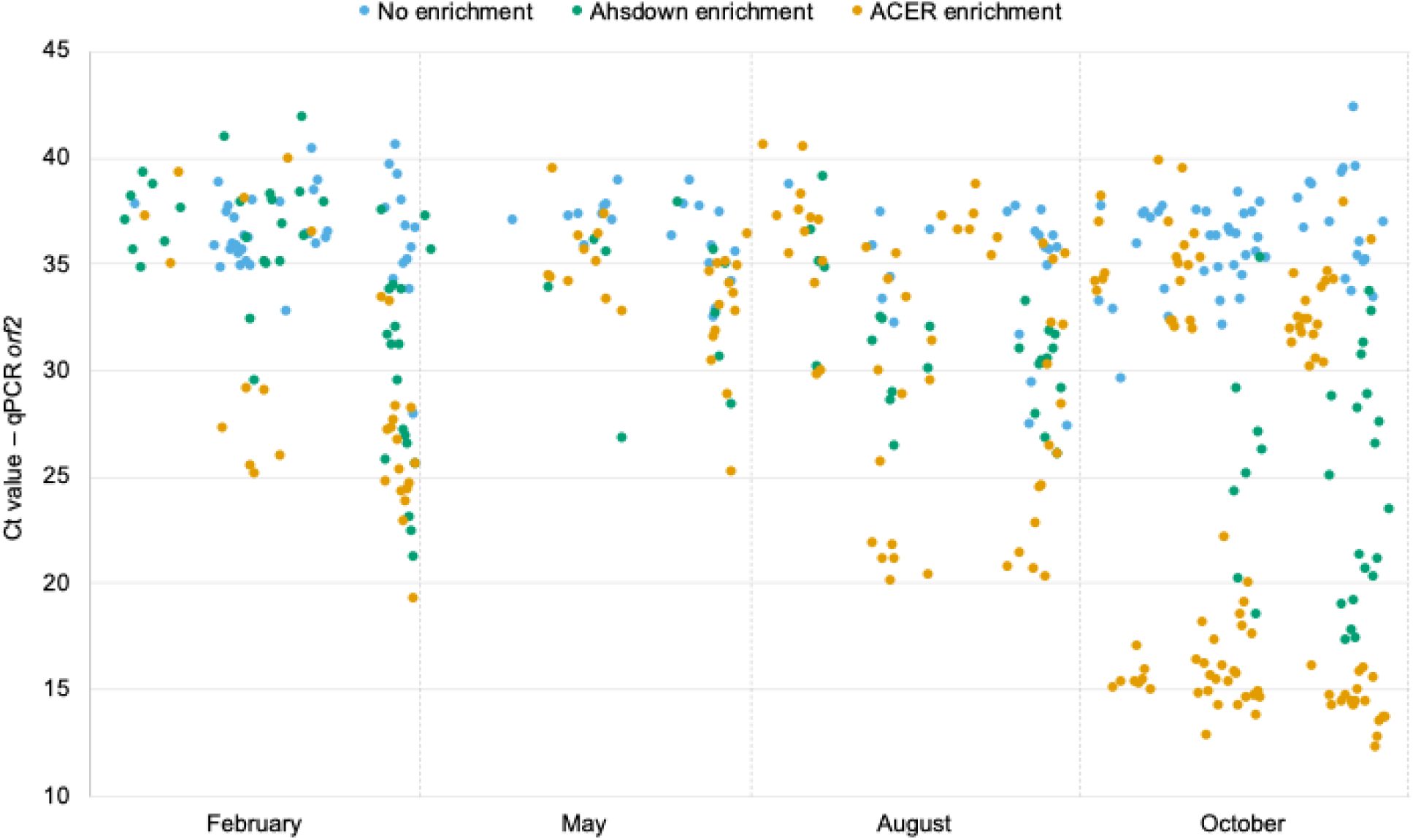
Sample-level comparison of Ct values obtained by qPCR targeting the TTS1-*orf2* gene of *B. pseudomallei* following no enrichment, Ashdown enrichment or ACER enrichment of soil samples collected in 2025. Each point represents an individual sample. Lower Ct values indicate higher amounts of target DNA. In October, 1 mL of Ashdown broth from sites 1 to 5 were not collected, hence those samples (n = 80) were not tested by qPCR.

Based on the 54 samples that tested qPCR-positive following all three pre-treatment protocols, Spearman’s rank correlation analysis was used to assess whether the relative ranking of samples according to qPCR Ct values ––a proxy of *B. pseudomallei* DNA abundance–– was consistent across pre-treatment protocols (Fig 5). A strong positive rank correlation was observed between Ashdown and ACER enrichments (ρ = 0.68, *p* < 0.001), indicating that samples tended to maintain a similar relative ranking following either enrichment protocol (Fig 5C). In contrast, no significant rank correlation was observed between no enrichment and Ashdown enrichment (ρ = 0.06, *p* = 0.693) or between no enrichment and ACER enrichment (ρ = 0.02, *p* = 0.858), indicating that the relative ranking of samples based on qPCR results without enrichment was not preserved following either enrichment protocol.

**Fig 5.**
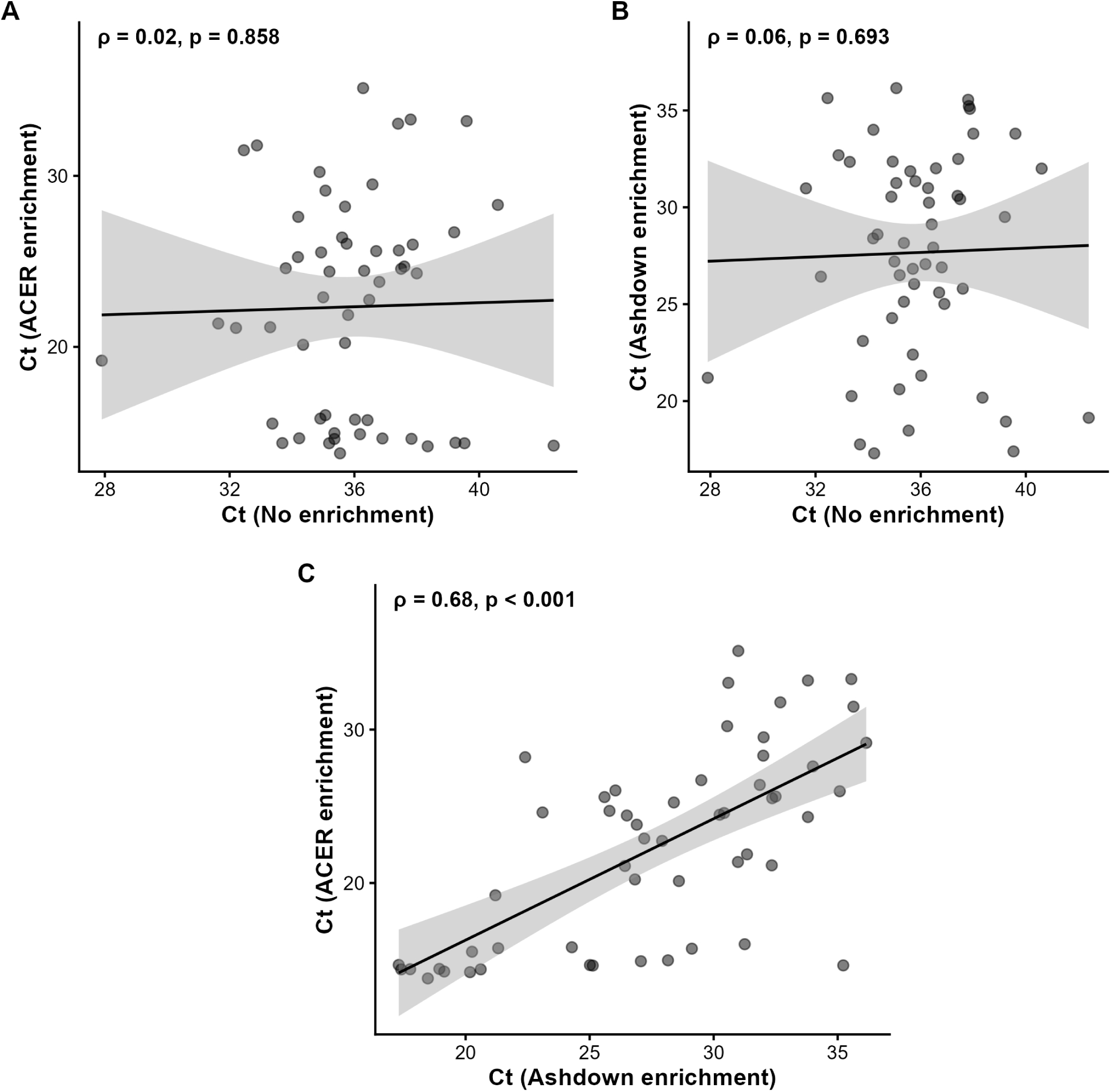
Scatterplots comparing Ct values between pre-treatment protocols for the 54 samples that tested qPCR-positive following all three protocols. (A) No enrichment vs. ACER enrichment; (B) No enrichment vs. Ashdown enrichment; (C) Ashdown enrichment vs. ACER enrichment. Black lines represent linear regression fits, with shaded areas indicating 95% confidence intervals. Rho (ρ) and *p*-values from Spearman’s rank correlation analysis are shown on each panel.

### Modeling of qPCR Ct values across three pre-treatment protocols

A GLMM was used to formally compare Ct values across the three protocols among the 54 samples that tested qPCR-positive following all three pre-treatment protocols, accounting for the paired structure of the data (i.e., the same samples tested under each protocol). The model-estimated mean Ct values were 36.7 for no enrichment, 28.4 for Ashdown enrichment, and 23.1 for ACER enrichment (Fig 6, Table S3). Pairwise comparisons showed that, compared with no enrichment, the mean Ct was reduced by 8.2 cycles with Ashdown enrichment and by 13.6 cycles with ACER enrichment, with both differences being significant (*p* < 0.0001). The reduction in Ct was therefore greater with ACER enrichment than with Ashdown enrichment, with a mean difference of 5.3 cycles between the two enrichment protocols (*p* < 0.0001).

**Fig 6.**
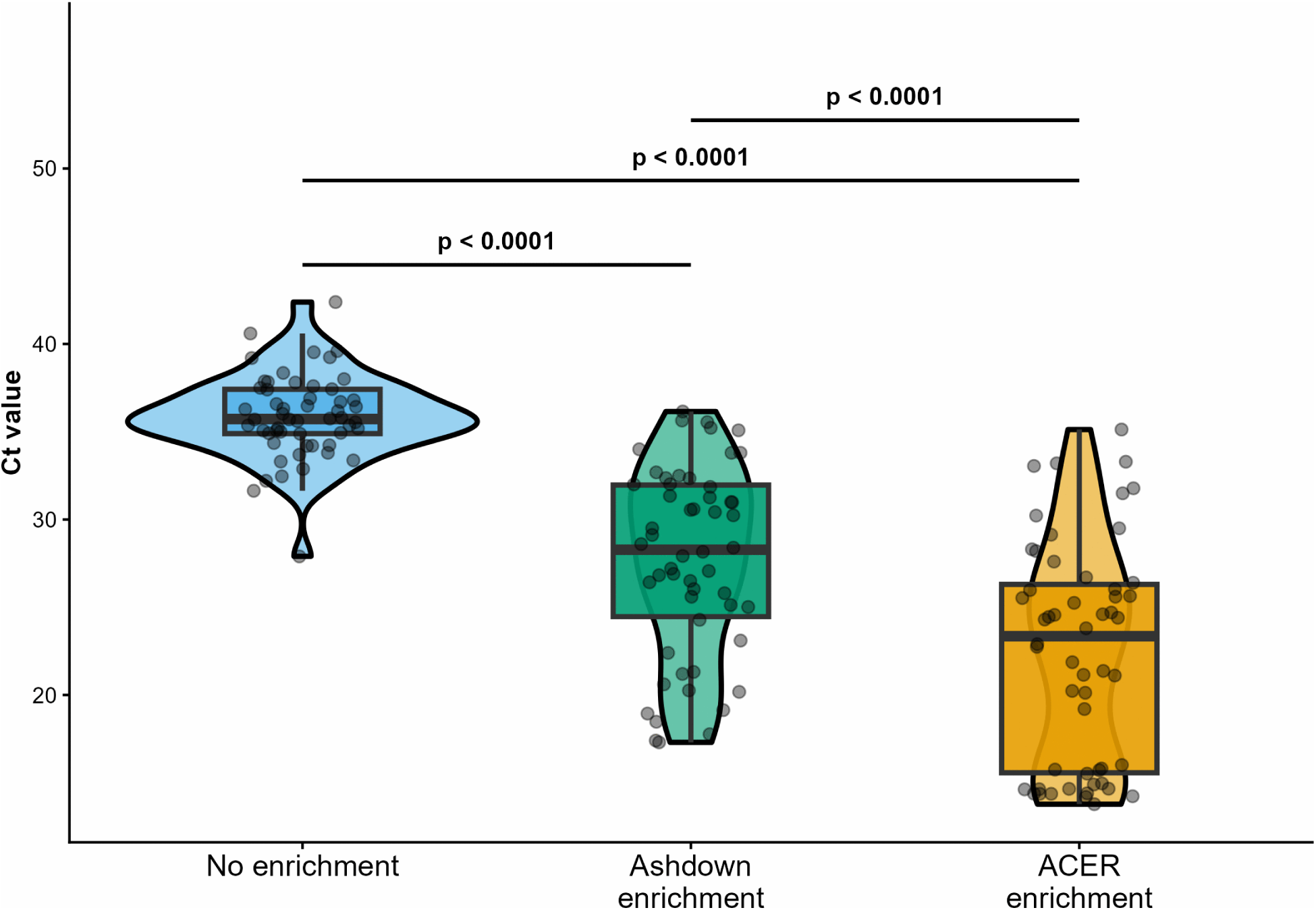
Distribution of Ct values obtained by qPCR targeting the TTS1-*orf2* gene of *B. pseudomallei* following no enrichment, Ashdown enrichment or ACER enrichment of soil samples. Lower Ct values indicate higher amounts of target DNA. Violin plots represent Ct value distributions, individual points represent samples, and boxplots indicate medians and interquartile ranges. Brackets indicate Tukey-adjusted pairwise comparisons (*p*-values).

### Sensitivity analysis of all results

Excluding February, during which a different supernatant-to-broth ratio was used for ACER enrichment, yielded slightly different results than with the full dataset.

For qPCR, ACER enrichment remained associated with higher odds of detection than no enrichment (OR = 3.32, 95% CI = 2.01–5.49 *vs* OR = 1.87 in the full dataset) and than Ashdown enrichment (OR = 3.79, 95% CI = 2.27–6.31 *vs* OR = 2.06 in the full dataset). In pairwise comparisons of Ct values, compared with no enrichment, the mean Ct was reduced by 8.7 cycles with Ashdown enrichment (*vs* 8.2 in the full dataset) and by 14.9 cycles with ACER enrichment (*vs* 13.6 in the full dataset), with both differences being significant (*p* < 0.0001).

For culture, isolates were recovered following no enrichment only in February, precluding a quantitative comparison with the “no enrichment” protocol when excluding this sampling period. ACER enrichment still yielded higher odds of detection than Ashdown enrichment (OR = 3.70, 95% CI = 1.32–10.38), although this association was attenuated compared with the full dataset (OR = 6.17).

Sensitivity analysis results are summarized in S4 Table, which includes the results described above as well as comparisons for the other analyses conducted on the full dataset.

## Discussion

As melioidosis cases are increasingly reported in countries beyond Southeast Asia and tropical Australia, reliable tools to confirm true endemicity, i.e., the local environmental presence of *B. pseudomallei,* are increasingly needed. Environmental surveys around melioidosis cases can help provide clues about transmission pathways or confirm potential sources of infection. In the Northern Territory of Australia, several clonal clusters of melioidosis have been linked to contamination through community water supplies [23,24]. Similarly, in Ubon Ratchathani province in northeastern Thailand, two melioidosis patients were infected with a *B. pseudomallei* genotype that was also present in their drinking water [25]. More recently, environmental surveys have been key to confirming that *B. pseudomallei* is locally established in historically non-endemic melioidosis countries, e.g., in the southern USA [26] and the Caribbean [5]. Environmental surveys are also increasingly being used to characterize the distribution of environmental *B. pseudomallei*, e.g., in Gabon [27], and to provide a better picture of *B. pseudomallei* diversity [28]. However, the challenges associated with detecting and isolating *B. pseudomallei* from soil have hampered efforts to characterize its environmental ecology, spatial distribution, and persistence.

The aim of our work was to evaluate and compare culture and molecular detection, with or without selective enrichment, for the detection of *B. pseudomallei* in naturally-contaminated soils collected over one year in Amnat Charoen province, northeastern Thailand. Our results showed a major discrepancy between bacterial isolation and molecular detection. Culture yielded only 3, 8 and 39 positive samples after no enrichment, Ashdown enrichment and ACER enrichment, respectively, whereas qPCR detected *B. pseudomallei* DNA in 133, 105 and 199 samples using the same three pre-treatment protocols.

ACER enrichment markedly increased the qPCR signal, resulting in lower Ct values than following no enrichment. Among the samples that tested qPCR-positive following all three pre-treatment protocols, the model-estimated mean Ct was 13.6 cycles lower after ACER enrichment than after no enrichment. Because a reduction of one Ct value corresponds approximately to a doubling of target DNA under optimal amplification conditions, this difference would correspond to a substantial increase in target DNA if amplification efficiency were comparable. However, this quantitative interpretation should be treated with caution after enrichment, as Ct values no longer solely reflect the initial bacterial concentration in soil but also incorporate differences in bacterial growth dynamics during enrichment incubation. Indeed, the absence of significant rank correlation between non-enriched samples and either Ashdown- or ACER-enriched samples indicates that the relative ranking of samples based on qPCR results without enrichment was not preserved following enrichment. Conversely, the strong rank correlation between Ashdown- and ACER-enriched samples suggests that both enrichment protocols produced similar relative rankings among samples after bacterial multiplication. Thus, enrichment substantially modifies the quantitative relationship between samples, potentially due to differences in bacterial growth dynamics, physiological state, or interactions with competing microorganisms present in the soil matrix.

Importantly, enrichment incubation periods differed between Ashdown (two days) and ACER (nine days), which may have contributed to the differences in bacterial loads observed after enrichment. A previous study compared TBSS-C50, TBSS-C50-based erythritol, and ACER broth enrichments following 2, 5, and 9 days of incubation [10]. Median colony counts were significantly higher following enrichment in ACER broth than in TBSS-C50 broth (*p* = 0.001), while TBSS-C50-based erythritol broth showed the lowest recovery at all time points. Ashdown enrichment was not included in that study. These findings suggest that both the enrichment medium and incubation duration may influence *B. pseudomallei* recovery; however, their relative contributions cannot be separated in our study as Ashdown and ACER protocols differed in both medium composition and incubation duration.

Soil was sampled during four periods over one year to assess the potential effect of sampling period on *B. pseudomallei* detection and isolation using different methods. Our results suggest that the additional detection provided by ACER enrichment over no enrichment was not consistent across sampling periods, whereas the higher detection associated with ACER compared with Ashdown was consistent throughout the year. Previous studies conducted in endemic regions have demonstrated increased environmental detection during periods of intense rainfall or typhoon activity, when soil disturbance and water redistribution may enhance bacterial dispersion and exposure risks [29]. In February and May, there was no statistically significant difference in paired detection rates between no enrichment and ACER enrichment, whereas the difference became statistically significant in August and October. This pattern may indicate that the benefit of enrichment is less pronounced when *B. pseudomallei* abundance in the original soil is very low. One possible explanation is stochastic variation during subsampling and pipetting. When *B. pseudomallei* abundance is very low, small differences in the amount of target bacteria transferred to different aliquots for different protocols may substantially affect detection or enrichment success. Conversely, at higher abundance, target bacteria may be more consistently represented across aliquots, reducing this stochastic effect. Our results suggest that the timing of environmental sampling may influence the probability of detecting *B. pseudomallei*. When abundance is predicted to be low, processing multiple soil replicates, broth aliquots, and/or agar plates may increase the likelihood of detecting *B. pseudomallei*, which may be present in only a small proportion of the sample.

The performance of culture or molecular detection following different pre-treatment protocols highlights an important distinction between bacterial isolation *versus* detection objectives. For a surveillance strategy aiming to characterize the environmental distribution or establish the presence of *B. pseudomallei,* ACER enrichment may be useful for increasing the likelihood of molecular detection probability and enhancing the qPCR signal. However, when the objective is to recover viable isolates for downstream applications, such as whole-genome sequencing, strain characterization, antimicrobial resistance studies, or virulence investigations, substantial limitations remain. In our study, despite the strong increase in molecular detection after ACER enrichment, only a fraction of qPCR-positive samples yielded isolates. This discrepancy was particularly evident in samples with lower initial bacterial abundance or during periods of reduced environmental contamination, where *B. pseudomallei* was likely outcompeted by abundant background microorganisms despite selective enrichment.

The difficulty of recovering environmental isolates is not specific to our study and discrepancies between culture isolation and molecular detection have been reported previously in other endemic regions. In Australia, 104 soil samples were randomly collected in Darwin rural area and the presence of *B. pseudomallei* was tested with both methods. The detection rate was 4% by culture and 12% by qPCR with soils collected 10 cm deep, and 15.2% by culture and 21.5% by qPCR with soils collected 40 cm deep [30]. In Laos, 100 soil samples were collected from a highly contaminated section of a rice paddy in the Vientiane Province, and the *B. pseudomallei* detection rate by qPCR post-enrichment was significantly higher (84%) than that by culture (44%) [31]. In Thailand, out of 40 soil samples collected in Ubon Ratchathani province, 26 samples were *B. pseudomallei* positive by both culture and qPCR, and seven samples were qPCR-positive but culture-negative [32]. In South West India, 517 out of 1,704 soil samples were positive by qPCR, whereas only two samples yielded culturable *B. pseudomallei* isolates [33]. These observations illustrate the higher detection rates often obtained with molecular methods compared with culture and highlight their value for mapping the environmental presence of the bacterium and identifying potential areas of increased exposure. Because isolation of environmental strains of *B. pseudomallei* remains a critical objective [12], future improvements should focus not only on increasing detection but also on enhancing the selective recovery of viable strains, particularly from soils with low *B. pseudomallei* loads and high microbial diversity. Combining optimized selective enrichment, targeted suppression of competing flora, and molecular screening strategies may provide a more efficient workflow for environmental surveillance and strain recovery.

This study has several limitations. First, our evaluation of culture-based methods was limited to a single protocol, i.e., plating on Ashdown agar followed by a 4-day incubation at 42°C. Other culture conditions might improve the recovery of *B. pseudomallei* isolates. Second, the supernatant-to-broth ratio used for ACER enrichment protocol differed between sampling periods (0.5 mL in 10 mL in February vs 1 mL in 9 mL otherwise). A more diluted supernatant would be expected to reduce, rather than inflate, the apparent sensitivity of ACER enrichment during that period, so including February should bias our findings conservatively. Indeed, our sensitivity analysis excluding February did not change the direction of any association. Finally, conducting the various protocols required 3.11 mL of soil supernatant: 1 mL for DNA extraction without enrichment, 1 mL for each enrichment broth, and 110 µL for direct plating on agar. As 3-mL microtubes were not available, two aliquots of soil supernatant were collected into two 2-mL microtubes before being distributed among the different protocols. In samples with low *B. pseudomallei* abundance, this subsampling step may have introduced variability between aliquots. For example, the bacterium may have been captured in the aliquot used for direct DNA extraction and qPCR but not, or at lower abundance, in the aliquot transferred to an enrichment broth.

Overall, our findings indicate higher detection rates for molecular methods than for culture when detecting *B. pseudomallei* in environmental soil samples. Furthermore, ACER enrichment increased molecular detection of environmental *B. pseudomallei* and produced lower Ct values than no enrichment or Ashdown enrichment, although its apparent benefit varied across sampling periods. ACER enrichment may therefore be useful for future environmental surveillance studies, while further work is needed to enhance the recovery of viable isolates.

## Acknowledgements

We thank the village chief of Kham Pha O for his help as well as all the private land owners who allowed access to their land for the soil sampling. We also thank the members of the Land Development Department (LDD) Amnat Charoen office and the villagers that contributed their help for the soil sampling. We thank the staff at the Department of Microbiology and Immunology, Faculty of Tropical Medicine, Mahidol University, for their kind support to this study. We are also grateful to Rathanin Seng from Mahidol University Amnat Charoen Campus, for his logistical support at the start of the project, and to Didtawa Suttisak for her help preparing ACER broth.

